# A Hierarchical Method to Analyze Protein-DNA Interfaces

**DOI:** 10.1101/2024.07.18.604047

**Authors:** Amol Tagad, G. Naresh Patwari

**Affiliations:** Department of Chemistry, Indian Institute of Technology Bombay, Powai, Mumbai 400076 INDIA

## Abstract

The accessibility of genetic information and packaging of long chromosomal DNA into micron-sized nuclei is a great example of protein-DNA interaction. A large number of protein-DNA structures are available in the database and is continuously increasing. The analysis of such huge data to extract meaningful insights, several computational tools such as hydrogen bonding, SASA, all intermolecular heavy atom-to-atom contacts, as well as web-server and databases such as DNAproDB, PDIdb, PDB2PQR, Bio-python are available. Generally, the interaction between the protein and DNA is analyzed based on all heavy atom-to-atom contact matrices which are computationally expensive. Herein, a new and robust hierarchical approach is developed to analyze the protein-DNA interface. The present hierarchical method comprises two steps; the first step is to recognize the protein residue-nucleotide pairs at the interface by employing pairwise distance cut-off between the Cα of the protein residues and 05′ of the nucleotide and the second step is to calculate heavy atom-to-atom contact matrix using the first step as a qualifier. This method reduces the computational cost by three orders of magnitude making it tractable even on personal computers. On the whole, the protein-DNA interface is dominated by the arginine residues with a notable presence of lysine, serine, and tyrosine, highlighting the pivotal role of electrostatic interactions and hydrogen bonding in aggregation.

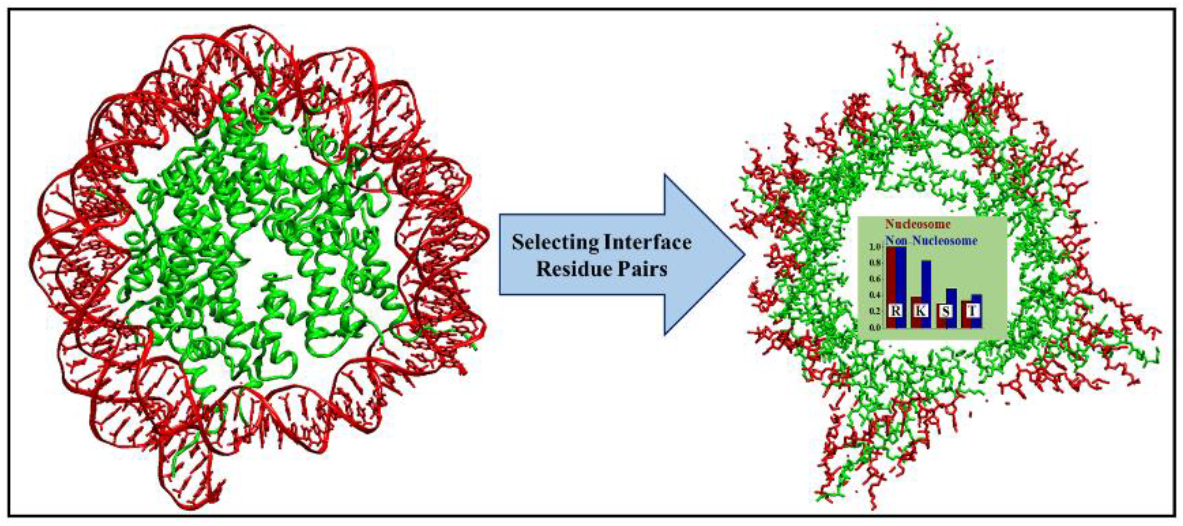

## INTRODUCTION

The interaction between protein and DNA is one of the most important processes as genetic information is accessible only when proteins engage with DNA. Further, several physiological processes including transcription, gene regulation, replication, DNA repair and modification involve protein-DNA interaction.^1−7^ On the other hand, the length of the chromosomal DNA is very long, hence has to be packed efficiently in a micron-sized nucleus, which is generally mediated by histones. Alternatively, the interaction between proteins and DNA in many cases is specific to local binding sites for their function and these interactions are crucial for healthy functioning and any abnormalities can result in a wide variety of diseases such as cancer, neurological disorders, diabetes, obesity and cardiovascular disease.^8−10^

Numerous experimental and computational methodologies are being used to investigate and understand the intricacies of protein-DNA(PD) interaction, and structure obtained using X-ray crystallography and nuclear magnetic resonance spectroscopy, provide valuable insights. In general, a DNA binding protein contains specific structural motifs such as helix-turn-helix(HTH), helix-loop helix (HLH),^11^ basic leucine zipper (bZIP),^12,13^ zinc-linker-helix,^14 −16^ and β sheets^17^ which specifically recognize and bind to the target DNA. The DNA binding motifs of proteins are target specific, with HTH/HLF or helix-loop-helix specifically binding to the major grooves,^12,15,18^ while zinc-linker-helix motif recognizes the base triplets,^19,20^ as in the case of CCG recognition in CBF3-DNA complex.^21,22^ Structural analysis reveals that the positively charged and polar residues (R, K, T and S) primarily interact with the DNA. This analysis has also revealed that the interaction between protein and DNA is predominantly (about ⅔) with the phosphate backbone (non-specific interaction) while the rest (about ⅓) constitute direct interaction of protein residues with the nucleobase (specific interaction).^23−25^ However, the intricate nature of interactions is influenced by the size, shape and sequence of the DNA.^26−28^ Several computational tools such as electrostatic energy of interacting residue-pairs,^29^ hydrogen bonding energy and patterns,^23,30^ solvent accessible surface area (SASA),^31^ binding free energy landscape^32^ are used to gain insights into the nature of protein-DNA interactions.

A large number of protein-DNA structures deposited in protein data bank (PDB) continues to increase.^33^ Several web-servers and databases are available to analyze the structural aspects of protein-DNA complexes such as DNAproDB,^34^ PDidb,^33^ PDB2PQR ^35,36^ Bio-python,^37^ PDA,^38^ OnTheFly.^39^ Generally, the between the protein and DNA is analyzed based on all heavy atom-to-atom contact matrices.^40−43^ However, in many cases the size of the protein-DNA is huge, therefore construction of all heavy atom-to-atom contact matrices becomes computationally expensive. In order to address this issue and render the analysis of protein-DNA computationally tractable, a new hierarchical approach is developed. This method is based on our earlier work on analyzing peptide aggregates.^44^ The newly developed method was used to analyze 1937 protein-DNA complexes, of which 258 structures were categorized as nucleosomes (keywords: Chromosome, NCP, Nucleosome) and the remaining 1679 structures correspond to protein-DNA complexes of transcription factors, DNA polymerases, DNA binding protein and other, in which a segment of DNA interacts with the protein (termed non-nucleosome in the present work). Figure 1 depicts representative structures of nucleosome and non-nucleosome protein-DNA complexes. The present work explores the quantitative correlation between protein-DNA interactions and identifies both specific and non-specific interactions in a residue pairwise fashion.

**Figure 1.**
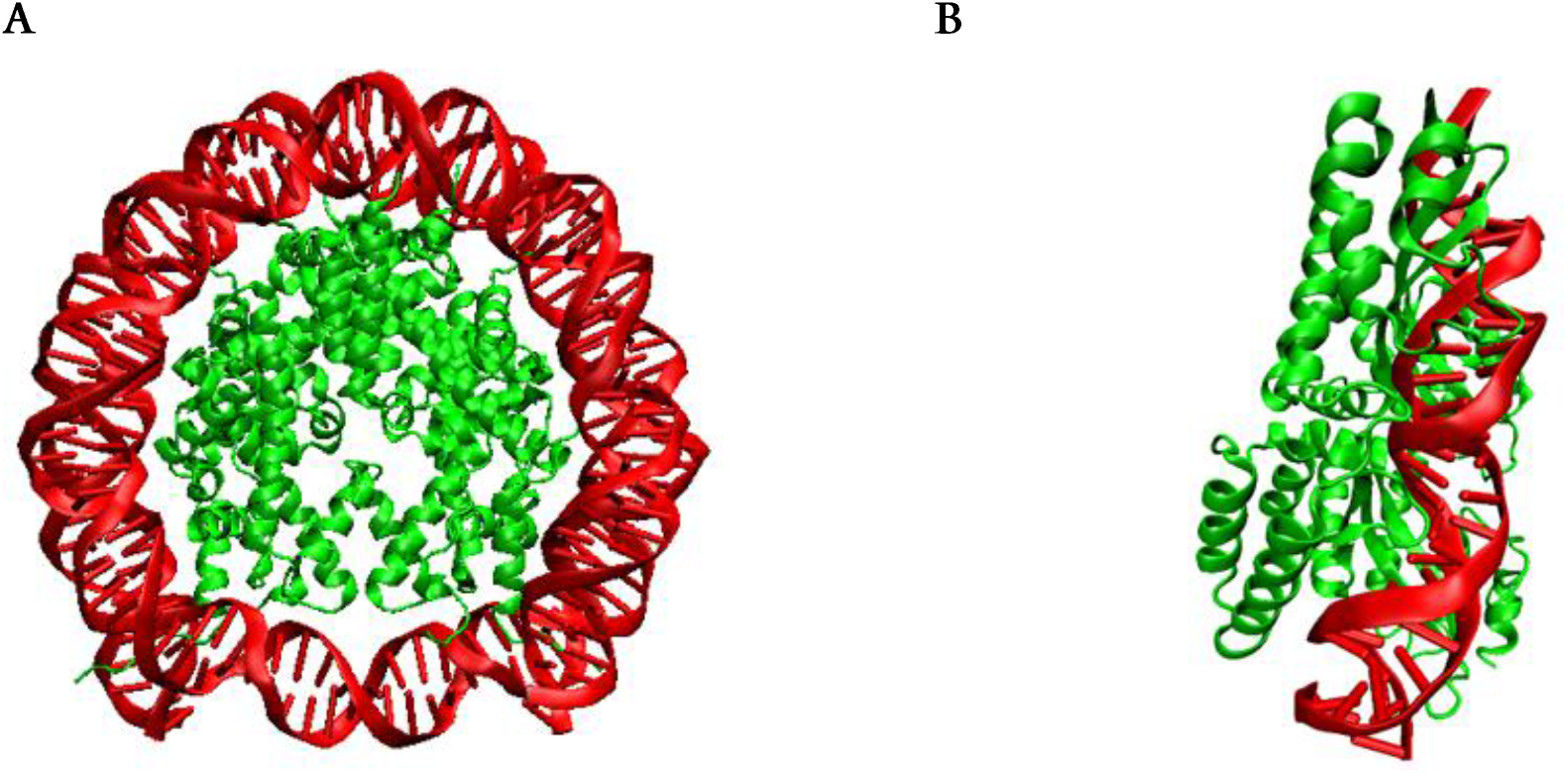
Representative structures of the DNA protein complex classified as (A) nucleosome (PDB-ID 1M19) and (B) non-nucleosome (PDB-ID: 2EX5). Both structures are represented in cartoon form wherein red and green colors represent DNA and protein, respectively.

## METHODOLOGY

To begin with 1937 structures of protein-DNA complexes deposited in the RCSB database,^45^ with a resolution better than 3.5 Å were downloaded (see Table SI for the PDB-ID list). Protein-DNA complexes involving single-stranded DNA and DNA sequences with fewer than 30 base pairs were not included in the dataset. The pairwise distances between Cα of protein residues and 05′ of nucleotides were calculated using GROMACS software package,^46^ which results in a *mxn* matrix, where *m* and *n* are the number of residues in protein and nucleotides in DNA,

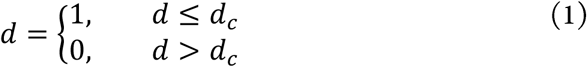

Cα−O5′ distances (*d*) were converted to binary index by imposing distance cut-off (*dc*) criteria. ^44^ Only those residues of protein with binary index **1** were considered for the interaction with DNA and others were neglected. This indexing allows sorting and selecting of the protein residues at the interface and neglecting the buried residues, which are not directly involved in the interaction with the DNA. Imposing the cut-off criterion, the pair-wise distances between the heavy atoms of interacting nucleotide-protein residue pairs were evaluated and interactions between the heavy atoms were constrained to 0.4 nm, for the final analysis. Setting the cut-off criteria on the pairwise distances between Cα of protein residues and 05′ of nucleotides leads to computationally efficiency over the construction of the entire heavy atom-to-atom contact matrix between all the nucleotide-protein pairs in the complex. The interaction between the nucleotides and the protein was further classified based on the sub-class of interactions of the heavy atoms and were categorized as (i) phosphate - protein, (ii) ribose - protein and (iii) nucleobase - protein and reciprocally as (iv) main chain - nucleotide and (v) side chain - nucleotide. Further, the hydrogen bonding interactions between the nucleotides and protein residues at the interface were calculated using the HBPLUS software,^47^ using two criteria, the first being the distance between hydrogen bond donor-acceptor heavy atoms being ≤ 0.35 nm and the second angle criteria ≥130°. The structures are visualized by VMD software.^48^

## RESULT AND DISCUSSIONS

This investigation is primarily focused on gaining insights into the interfacial interactions of protein DNA complex structures which are available in the PDB database. The primary objective is to identify the specific residue pairs present at this interface and to evaluate the nature of the interaction.^28^ Figure SI (see the Supporting information) depicts the relative population of all the amino acids within various cut-off distances of 0.80, 1.00 and 1.20 nm between the Cα atom of amino acid and the 05′ atom of nucleotide (Cα − 05′) for both nucleosome and non-nucleosome structures.^49^ On the other hand, Figure 2 shows the relative abundance of the various amino acids with 1.2 nm cut-off distance between Cα and 05′ for arginine and lysine and with 1.0 nm cut-off distance between Cα and 05′ for all the other amino acids, *vide infra*. This cut-off defines the interface between the DNA and the protein. In the case of nucleosomes, the positively charged amino acid arginine is more prevalent followed by lysine, which is expected since the DNA is negatively charged. The significant presence of polar side-chain amino acids such as serine and threonine in the case of nucleosomes, at the interface, suggests the importance of hydrogen bonding between protein and DNA. Surprisingly, apolar residues alanine, leucine, and isoleucine are abundant at the interface. In the case of non-nucleosomes, both the positively charged positively charged amino acids arginine and lysine have similar propensity. The distribution of other amino acids is quite widespread with a relatively low abundance of cysteine, methionine and tryptophan. Interestingly, even aspartate and glutamate have a significant presence at the interface.

**Figure 2.**
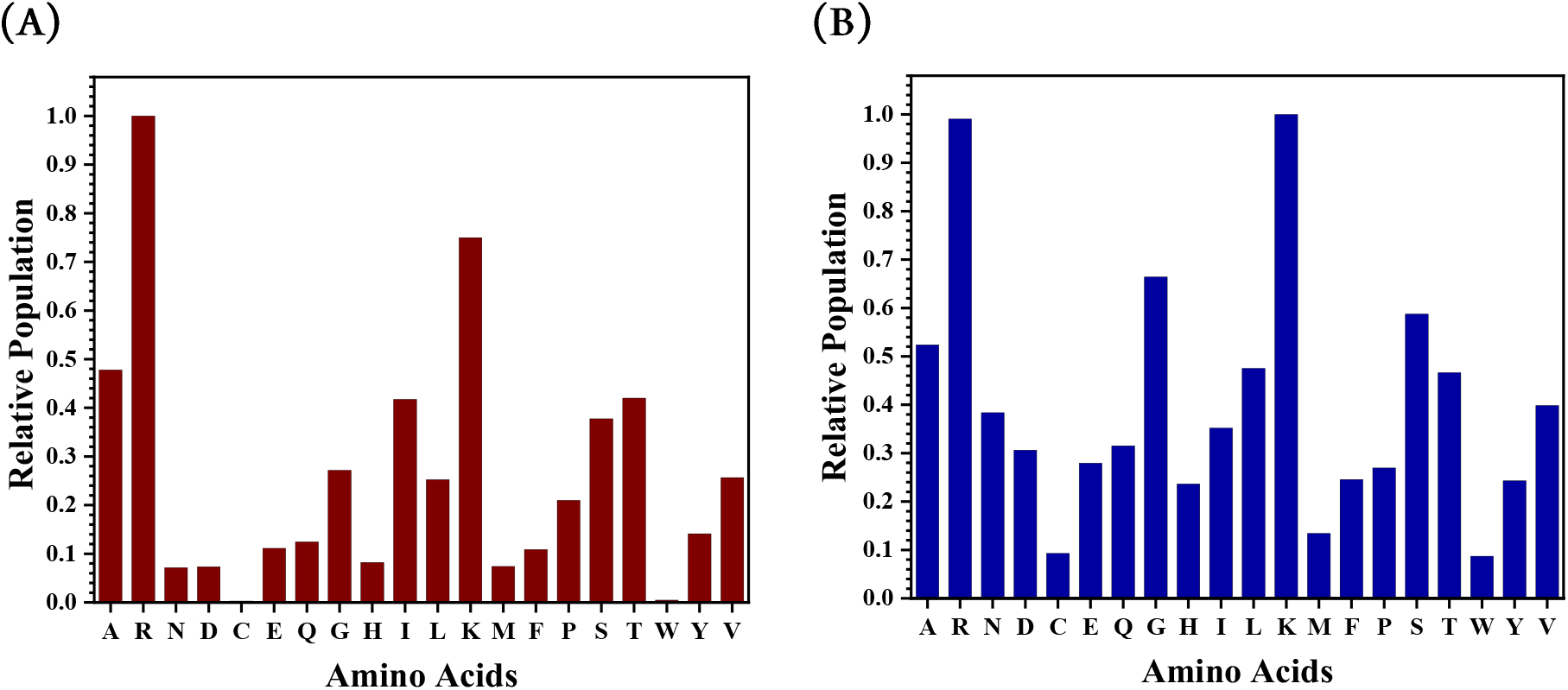
The relative population of all the 20 standard amino acids involved at the interface with 1.2 and 1.0 nm cut-off distances between Cα and 05′ for positively charged residues (arginine and lysine) and other residues, respectively for (A) nucleosomes and (B) non-nucleosomes. The number of PDB structures considered, with better than 3.5 Å resolution, for nucleosomes and non-nucleosomes were 258 and 1679, respectively.

To benchmark the new methodology Cα −O5′ distance cut-offs of 1.0 and 1.2 nm were considered for 10 nucleosome PDB structures (see Table S2 for the list), following which the heavy atom-to-atom contacts within cut-off distance of 0.40 nm between protein residues and the nucleotides were evaluated and the results are depicted in Figure 3. The results from the present hierarchical method (HM) were compared with the commonly used heavy atom (of all residues of protein) to heavy atom (of all nucleotides) methodology.^40−43^ Based on the relative population analysis, presented in Figure 3, it can be inferred that the Cα − O5′ distance cut-offs of 1.2 nm for arginine and lysine and 1.0 nm for all the other amino acids followed by 0.4 nm cut-off for the intermolecular heavy atom distances is adequate to capture details of the protein-DNA interaction at the interface.

**Figure 3.**
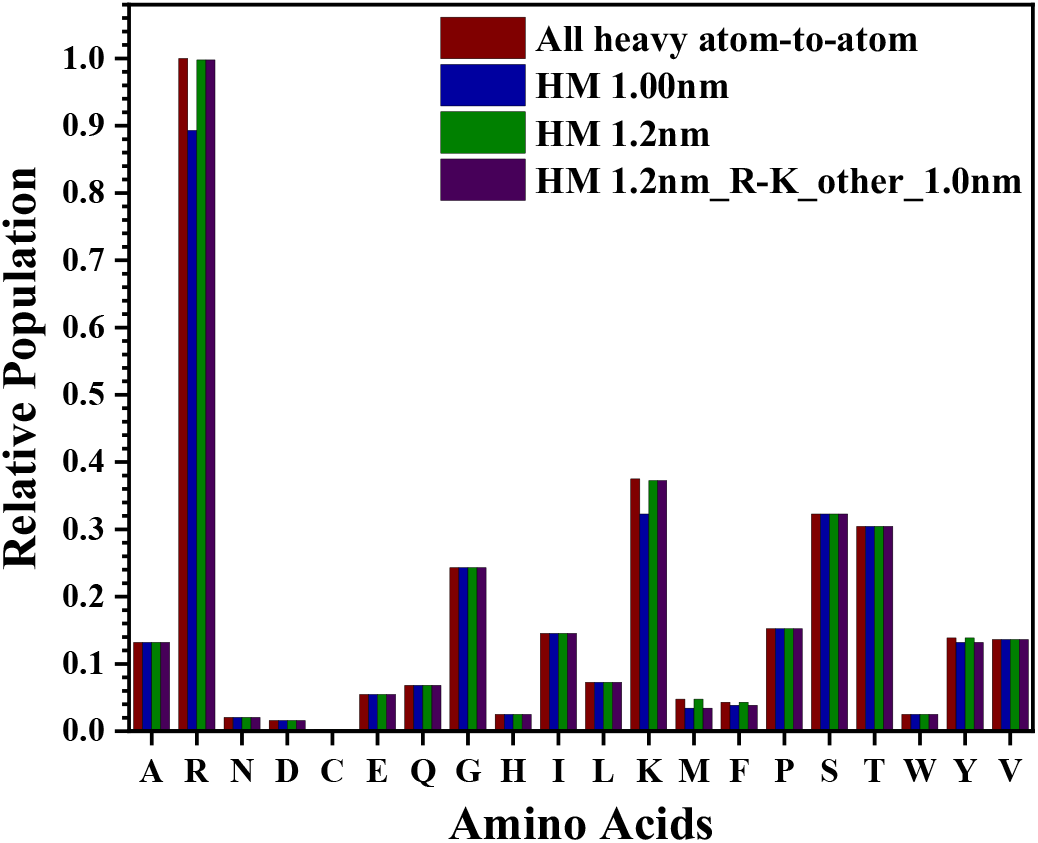
The relative population of all the 20 standard amino acids calculated using all the heavy atom-to-atom distances for representative 10 DNA-nucleosome PDB structures with the distance cut-off of 0.40 nm. Also shown are the relative populations using the hierarchical method (HM) of determining the protein residues with Cα − 05′ distance cut-off of 1.0 and 1.2 nm at the interface and calculating the heavy atom-to-atom distances for the sub-population at the interface with distance cut-off of0.40 nm. Notice that the Cα − O5′ distance cut-off of 1.2 nm for arginine and lysine and 1.0 nm cut-off for all the other amino acids is optimal.

It is essential to accentuate the significant difference in computational cost between the commonly used heavy atom (of all residues of protein) to heavy atom (of all nucleotides) method with the present hierarchical method (HM). For the 10 PDB nucleosome structures that were used to benchmark, the approximate time to generate all heavy atom-to-atom distance data using a using as single the Intel Xeon 8 core E3 processor took approximately 37,080 minutes. In contrast, the new HM method achieved the same analysis much more efficiently, requiring only 97 minutes (see Table S3 for the computation time requirements). Consequently, employing the HM method with 0.4 nm intermolecular heavy atom cut-off distance constrained with the Cα − O5′ cut-off distances of 1.2 nm for arginine and lysine and 1.0 nm for and for all other protein residue is an efficient protocol to evaluate the protein-DNA interface.

Once the cut-off criteria for Cα − 05′ is set, the population of the protein residues at the interface with DNA was estimated using intermolecular heavy atom distances with a cut-off of 0.4 nm. Figure 4 depicts the population protein residues at the interface for both nucleosome and non-nucleosome covering 258 and 1679 protein-DNA structures from the PDB. In the case of nucleosomes, arginine has the maximum population followed by lysine, threonine and serine in that order. In the case of nucleosomes, the population of arginine is significantly higher than lysine, in contrast to non-nucleosomes wherein the difference in the population of arginine and lysine residues is marginal. Further, in non-nucleosomes, unlike nucleosomes, the population distribution of other amino acids is quite widespread with a relatively low abundance of cysteine, methionine and tryptophan. Interestingly, even aspartate and glutamate have a significant presence at the interface.

**Figure 4.**
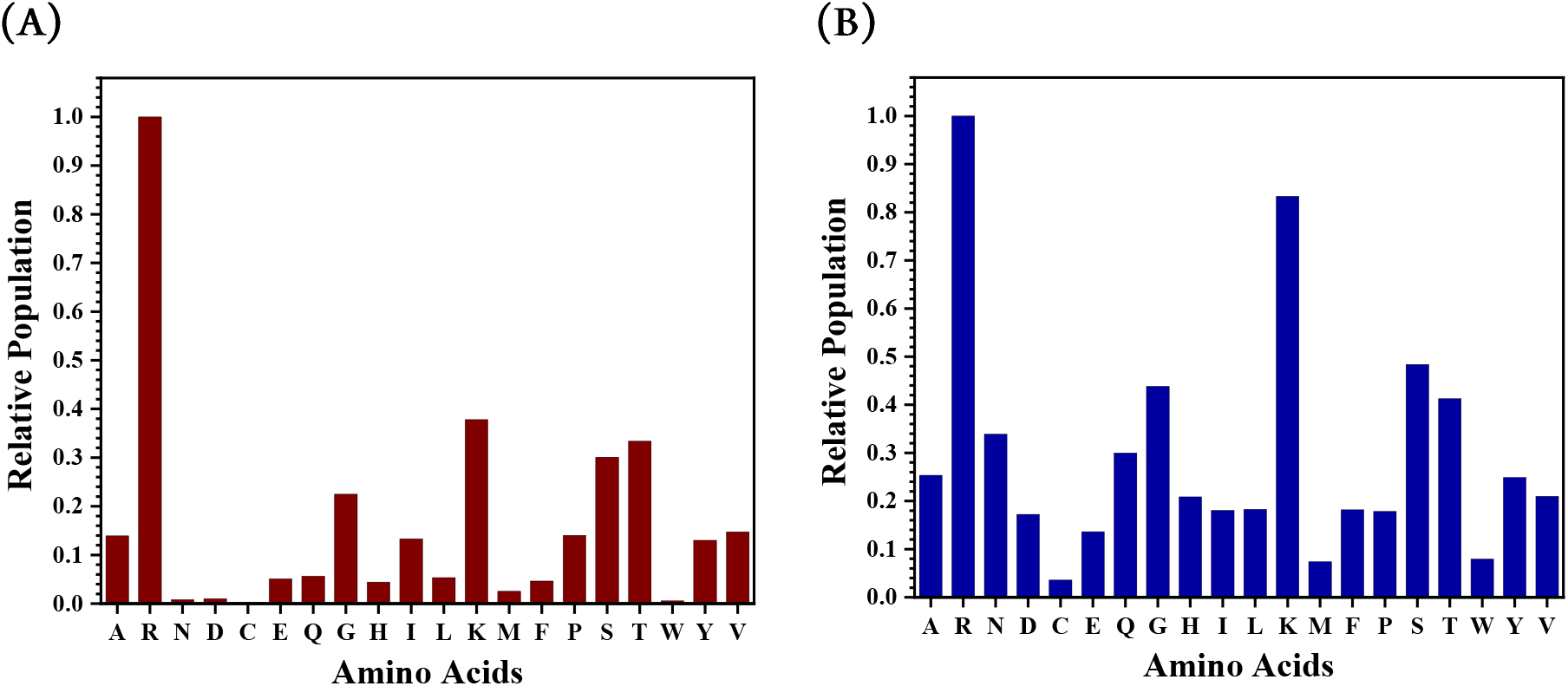
The relative populations of **(A)** nucleosomes and **(B)** non-nucleosomes using the hierarchical method (HM) of determining all the 20 standard amino acids with Cα − O5′ distance cut-off of 1.0 and 1.2 nm at the interface and calculating the heavy atom-to-atom distances for the sub-population at the interface with distance cut-off of 0.40 nm.

For a comprehensive understanding of the protein-DNA interface interactions, the HM method was further utilized to analyze the specificity of the interaction depending on the identity of the atom pairs involved and are categorized as the interaction of protein residues with (i) nucleobases, (ii) the ribose and (iii) the phosphate. On the other hand, the interaction of the nucleotides with the protein has two classes (i) main chain and (ii) side chain. Figure 5 depicts the statistics of the specific interactions. In general, the protein residues predominantly interact with the phosphate group, both for nucleosomes and non-nucleosomes. Enhanced interaction of amino acid residues with the ribose and nucleobases is observed in non-nucleosomes in comparison with nucleosomes, which is attributed to the sequence specificity of the protein-DNA interaction in non-nucleosomes. Similarly, the interaction of the side chain of the amino acid residues with the nucleotides is observed in non-nucleosomes which once again can be attributed to the specificity of the protein-DNA interaction.

**Figure 5.**
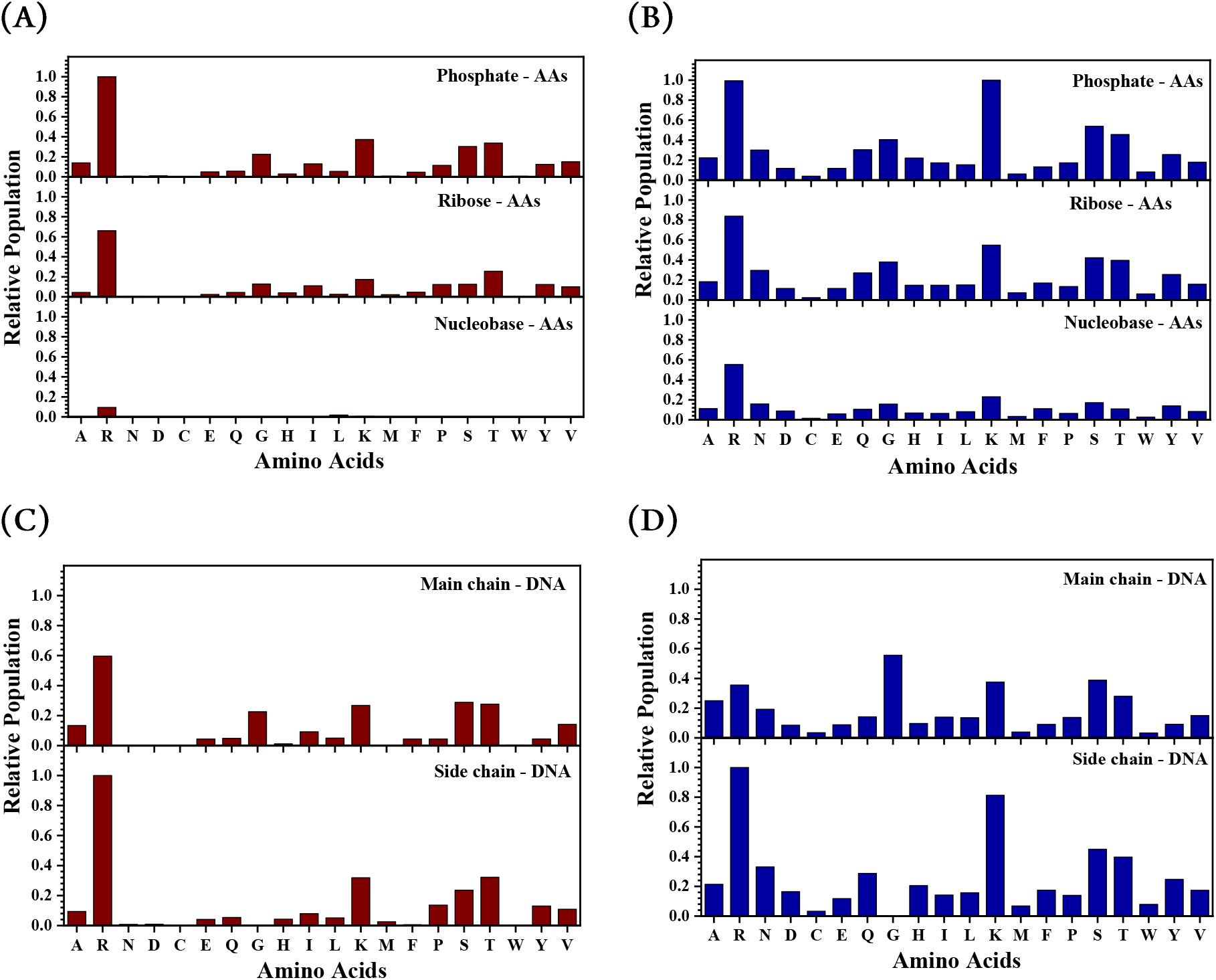
The Relative population plot of all standard 20 amino acids with DNA constituents **(A, B)** such as phosphate, ribose, and nucleobase, similarly protein constituent such as Main and Side chain with DNA (**C, D)** for both nucleosome and non-nucleosome within 0.40nm cut-off of intermolecular heavy atom contact.

In order to evaluate the role of hydrogen bonding at the protein-DNA interface, the hydrogen bonding propensities at the residue level for both nucleosomes and non-nucleosomes were evaluated using HBPLUS^47^ and are compared with the present HM approach and the results are presented in Figure 6. In general, the hydrogen-bonded contacts underestimate the interfacial interaction between the protein and DNA. Specifically, the residues arginine, lysine, serine and threonine have higher propensity to form hydrogen bonds in non-nucleosomes (72%) in comparison with nucleosomes (62%), which once again can be attributed to the specificity of the protein-DNA interaction in non-nucleosomes.

**Figure 6.**
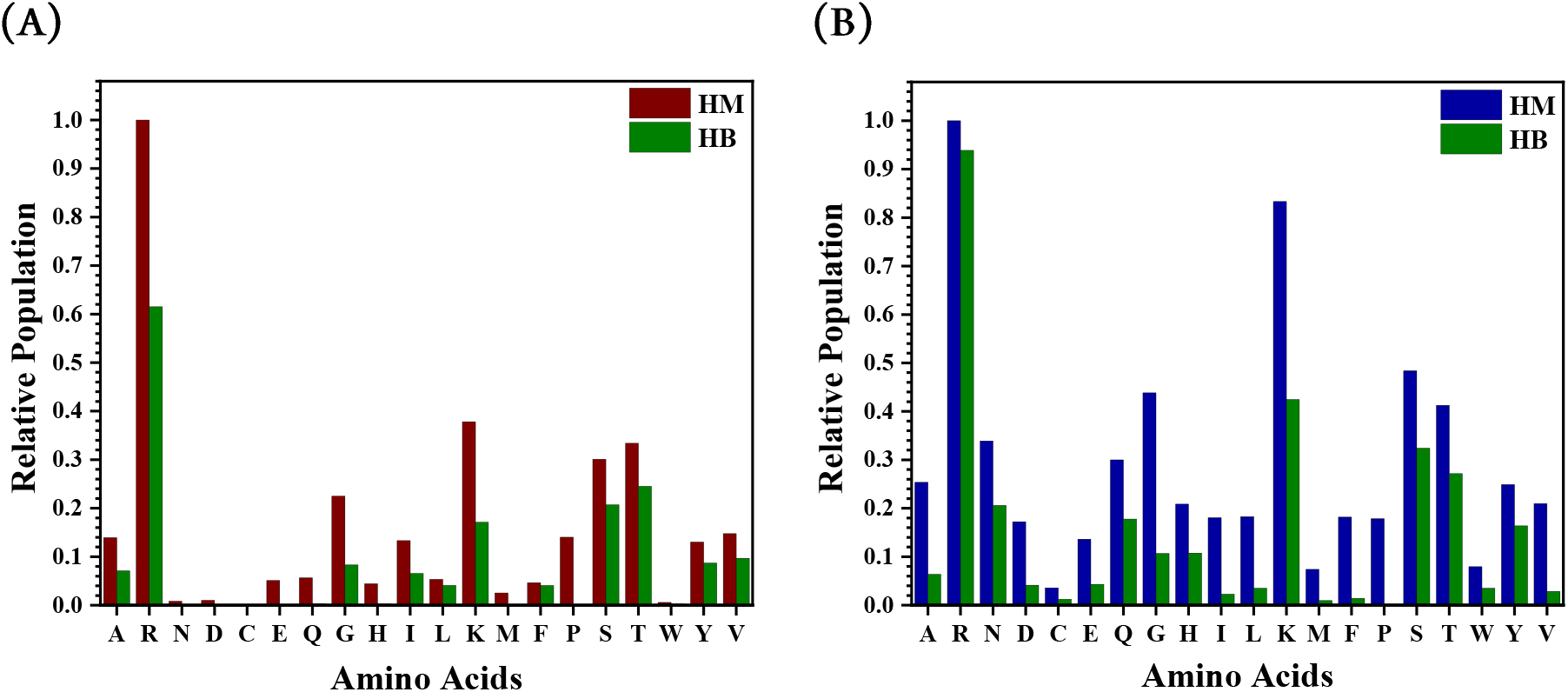
The relative population of all standard 20 amino acids by using the hierarchical method (HM) and hydrogen bonding (HB) for **(A)** nucleosome and **(B)** non-nucleosome. Note that the propensity for hydrogen bonding of arginine, lysine, serine and threonine is higher for non-nucleosomes (72%) over nucleosomes (62%)

## CONCLUSIONS

Herein, a hierarchical method is developed to investigate interfacial interactions in protein-DNA complexes reported in the PDB and a dataset consisting of 1,937 structures (258 nucleosomes and 1,679 non-nucleosomes) was analyzed. The hierarchical method computes the interface residue pairs by estimating the distance between the Cα atom of amino acid and the 05′ atom of nucleotide and selecting only the interfacial amino acid residues with a cutoff distance of 1.2 nm for arginine and lysine and 1.0 nm for the rest. A nested evaluation of all heavy atom-to-atom distances (0.4 nm) within the Cα − O5′ cut-off which defines the interface. This approach reduces computational costs significantly and this is tractable even on personal computers. The heavy atom-to-atom distances of interface residue pairs were then used to examine interactions between specific components of proteins (main and side chains) and nucleotides (phosphate group, ribose, and nucleobase). This analysis suggests that the nucleosomes have a greater arginine enrichment than non-nucleosomes. There is a greater proclivity for interactions with the phosphate and sugar moieties for the nucleosomes. On the other hand, interactions with nucleobases are sizable for non-nucleosomes, which is attributed to the sequence specificity of the interaction. In general, the protein-DNA interface is dominated by arginine, with substantial populations of lysine, serine and tyrosine.

## DATA AND SOFTWARE AVAILABILITY

The bash-script used for downloading the requisite PDB files from the RCSB Protein Data Bank, the implementation of aggregation analysis (see the flow chart in SI) and the data generated for benchmarking is provided at https://doi.org/10.5281/zenodo.10473357

## SUPPORTING INFORMATION

Flowchart to analyze the protein-DNA interface, list of PDB IDs used for analysis, table with computational time required for benchmarking and plot depicting population of the various amino acids at the interface (PDF).

## ACKNOWLEDGEMENTS

AT thanks the University Grants Commission (UGC) for the research fellowship. The authors wish to thank Dr. Reman Kumar Singh for his comments and suggestions. Authors gratefully acknowledge SpaceTime-2 supercomputing facility at IIT Bombay for the computing time. The support and the resources provided by ‘PARAM Brahma Facility’ under the National Supercomputing Mission, Government of India at the Indian Institute of Science Education and Research (IISER) Pune are gratefully acknowledged. This study is based upon a work supported in part by the Science and Engineering Research Board of the Department of Science and Technology (Grant no. CRG/2022/005470) and the Board of Research in Nuclear Sciences (BRNS Grant no. 58/14/18/2020) to GNP.

