## Supplementary material for "A Hierarchical Method to Analyze Protein-DNA Interfaces": Suplimentarly Information

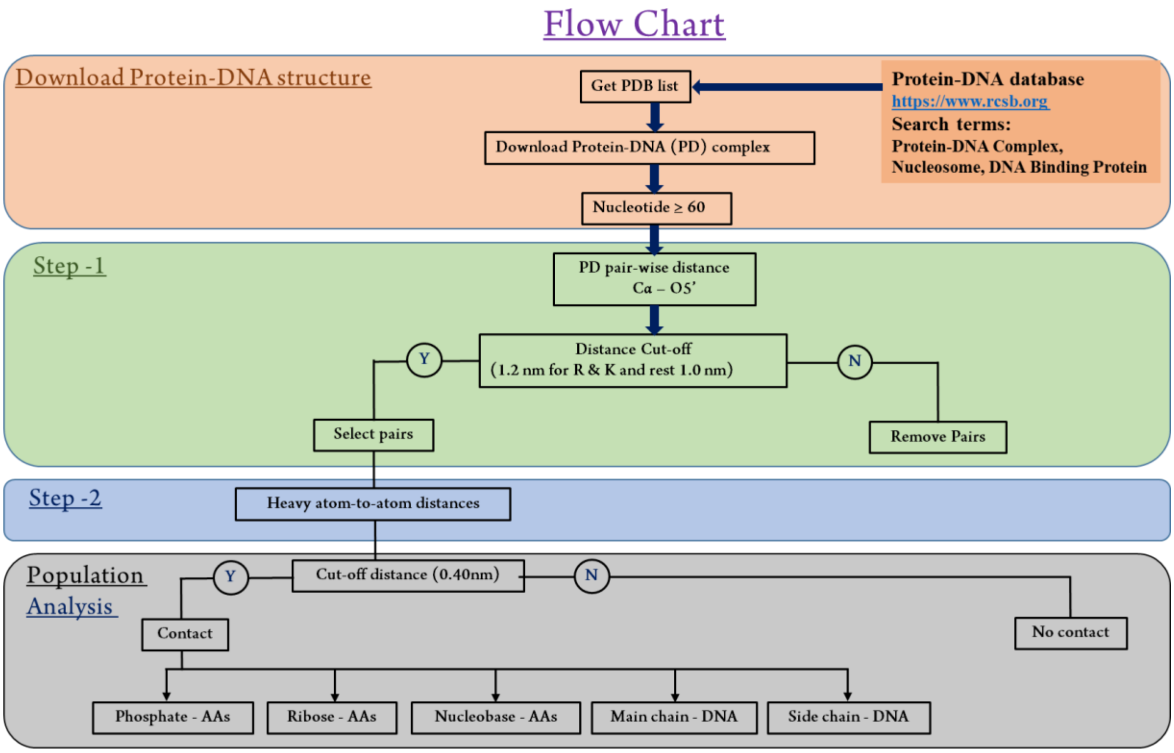

**Table S1.** The protein-DNA complex classified in nucleosome and non-nucleosome with a PDB ID.

| Type | PDB ID | Total |
|---|---|---|
| Nucleosome | 1AOI 1EQZ 1F66 1ID3 1KX3 1KX4 1KX5 1M18 1M19 1M1A 1P34 1P3A 1P3B 1P3F 1P3G 1P3I 1P3K 1P3L 1P3M 1P3O 1P3P 1S32 1U35 1ZLA 2CV5 2F8N 2FJ7 2NQB 2NZD 2PYO 3A6N 3AFA 3AV1 3AV2 3AYW 3AZE 3AZF 3AZG 3AZH 3AZI 3AZJ 3AZK 3AZL 3AZM 3AZN 3B6F 3B6G 3C1B 3C1C 3KUY 3KWQ 3KXB 3LEL 3LJA 3LZ0 3LZ1 3MGP 3MGQ 3MGR 3MGS 3MNN 3MVD 3O62 3REH 3REI 3REJ 3REK 3REL 3TU4 3UT9 3UTA 3UTB 3W96 3W97 3W98 3W99 3WA9 3WAA 3WKJ 3WTP 3X1S 3X1T 3X1U 3X1V 4INM 4J8U 4J8V 4J8W 4J8X 4JJN 4KGC 4KUD 4LD9 4QLC 4R8P 4WU8 4WU9 4X23 4XUJ 4XZQ 4YS3 4Z5T 4Z66 5AV5 5AV6 5AV8 5AV9 5AVB 5AVC 5AY8 5B0Y 5B0Z 5B1L 5B1M 5B2I 5B2J 5B31 5B32 5B33 5B40 5CP6 5CPI 5CPJ 5CPK 5DNM 5DNN 5E5A 5F99 5GSE 5GSU 5GT0 5GT3 5GTC 5GXQ 5JRG 5MLU 5OMX 5ONG 5ONW 5X7X 5XF3 5XF4 5XF5 5XF6 5XM0 5XM1 5Y0C 5Y0D 5Z23 5Z30 5ZBX 6DZT 6E0C 6E0P 6IPU 6IQ4 6IRO 6IY2 6JOU 6JR0 6JR1 6JXD 6JYL 6K1I 6K1J 6K1K 6KE9 6KIU 6KVD 6L9H 6L9Z 6LA8 6LAB 6LE9 6LER 6M4G 6MUP 6NJ9 6O1D 6O96 6OM3 6PA7 6PX1 6R94 6RYR 6S01 6SE6 6SEG 6T79 6T90 6T93 6UH5 6UPH 6V2K 6VEN 6WKR 6WZ5 6WZ9 6X59 6Y5E 6YOV 6ZHX 6ZHY 7A08 7COW 7CRP 7CRQ 7CRR 7D1Z 7D20 7E8D 7E8I 7EA5 7EA8 7EG6 7ENN 7JO9 7JOA 7K5X 7K5Y 7K60 7K61 7K63 7K6P 7K6Q 7K78 7KBD 7KBE 7KTQ 7LV8 7LYA 7LYB 7LYC 7NKX 7NKY 7NL0 7OH9 7OHA 7OHB 7OHC 7ON1 7PII 7R5R 7R5S 7SCZ 7SWY 7TAN 7TN2 7U46 7VBM 7VDT 7VDV 7XCR 7XCT 7XD0 7XD1 7XPX 7Y7I | 258 |
| Non-nucleosome | 10MH 1A02 1A1F 1A1G 1A1H 1A1I 1A1J 1A1K 1A1L 1A31 1A35 1A36 1A3Q 1A6Y 1AAY 1AIS 1AKH 1AM9 1AN2 1APL 1AU7 1AV6 1AZ0 1AZQ 1B3T 1B72 1B8I 1BC7 1BDH 1BDI 1BDT 1BDV 1BF4 1BF5 1BG1 1BPX 1BPY 1BPZ 1BRN 1BY4 1C8C 1C9B 1CA5 1CA6 1CBV 1CDW 1CGM 1CGP 1CIT 1CKQ 1CL8 1CLQ 1CMA 1CQT 1CW0 1CYQ 1CZ0 1D02 1D0E 1D1U 1D2I 1D3U 1D5Y 1D66 1DC1 1DDN 1DFM 1DGC 1DH3 1DIZ 1DMU 1DNK 1DP7 1DRG 1DSZ 1DU0 1EA4 1EBM 1ECR 1EFA 1EGW 1EHL 1EJ9 1EMH 1EMJ 1EO3 1EO4 1EOO 1EOP 1ERI 1ESG 1EVW 1EXI 1EXJ 1EYG 1EYU 1F0O 1F2I 1F44 1F4K 1F4R 1F6O 1FIU 1FJL 1FJX 1FLO 1FN7 1FOS 1FYL 1FYM 1G2D 1G2F 1GA5 1GD2 1GDT 1GLU 1GT0 1GTW 1GU4 1GU5 1GXP 1H0M 1H88 1H89 1H8A 1H9T 1HBX 1HCQ 1HCR 1HDD 1HJB 1HJC 1HLO 1HLV 1HLZ 1HUO 1HUZ 1HW2 1I3J 1I6J 1I7D 1I8M 1IAW 1IGN 1IHF 1IJS 1IJW 1IMH 1IO4 1IU3 1IXY 1J1V 1J3E 1J59 1J5O 1J75 1JB7 1JE8 1JEY 1JFI 1JFS 1JFT 1JGG 1JH9 1JJ6 1JJ8 1JK1 1JK2 1JKO 1JKP 1JKQ 1JKR 1JMC 1JNM 1JX4 1JXL 1K4S 1K4T 1K61 1K6O 1K78 1K79 1K82 1K8G 1KB2 1KB4 1KB6 1KBU 1KC6 1KEG 1KFS 1KIX 1KLN 1KRP 1KSP 1KU7 1L3L 1L3S 1L3T 1L3U 1L3V 1L5U 1LB2 1LE5 1LE8 1LE9 1LEI 1LLI 1LLM 1LMB 1LPQ 1LRR 1LWS 1LWT 1LWV 1LWW 1LWY 1M0E 1M3H 1M5R 1M5X 1MA7 1MDM 1MDY 1MEY 1MHT 1MJE 1MM8 1MNM 1MUH 1MUS 1MVM 1N3E 1N3F 1N48 1N56 1N6J 1NGM 1NH2 1NH3 1NJW 1NJX 1NJY 1NJZ 1NK0 1NK4 1NK5 1NK6 1NK7 1NK8 1NK9 1NKB 1NKC 1NKE 1NKP 1NOP 1NOY 1NVP 1NWQ 1O3Q 1O3R 1O3S 1O3T 1OCT 1OMH 1ORN 1ORP 1OTC 1OUZ 1OWF 1OWG 1OWR 1OZJ 1P47 1P59 1P7D 1P7H 1PA6 1PAR 1PDN 1PER 1PFI 1PH1 1PH2 1PH3 1PH4 1PH5 1PH6 1PH7 1PH8 1PH9 1PHJ 1PNR 1PT3 1PUE 1PUF 1PVI 1PVP 1PVQ 1PVR 1PYI 1PZU 1Q0T 1Q3U 1Q3V 1QAI 1QAJ 1QLN 1QP0 1QP4 1QP7 1QP9 1QPS 1QPZ 1QQA 1QQB 1QRV 1QSS 1QSY 1QTM 1QX0 1QZG 1QZH 1R49 1R4I 1R4O 1R4R 1R71 1R7M 1R8D 1R8E 1RAM 1RBJ 1RCN 1RFF 1RFI 1RG1 1RG2 1RGT 1RGU 1RH0 1RH6 1RM1 1RMV 1RPE 1RR8 1RRJ 1RRQ 1RRS 1RTA 1RUN 1RUO 1RVA 1RVB 1RVC 1RXV 1RZ9 1RZR 1S6M 1S9K 1SA3 1SC7 1SEU 1SFU 1SKM 1SRS 1SSP 1SUZ 1SX5 1T2K 1T2T 1T3N 1T8E 1T8I 1T9I 1T9J 1TAU 1TC3 1TGH 1TL8 1TQE 1TRO 1TRR 1TSR 1TTU 1TUP 1TX3 1U0C 1U0D 1U3E 1U45 1U47 1U48 1U49 1U4B 1U78 1U8B 1UA0 1UA1 1UBD 1UUT 1VAS 1VKX 1VOL 1VPW 1VRL 1VTL 1VTN 1VTO 1W0U 1WD0 1WD1 1WET 1WNE 1WTO 1WTP 1WTQ 1WTR 1WTV 1WTW 1WTX 1WVL 1X9N 1XC8 1XPX 1XSD 1XSL 1XSN 1XSP 1XYI 1YF3 1YFH 1YFI 1YFJ 1YFL 1YNW 1YO5 1YRN 1YSA 1YTB 1YTF 1Z19 1Z63 1ZAA 1ZET 1ZG1 1ZG5 1ZLK 1ZM5 1ZNS 1ZQA 1ZQB 1ZQC 1ZQD 1ZQF 1ZQG 1ZQH 1ZQI 1ZQJ 1ZQK 1ZQL 1ZQM 1ZQN 1ZQO 1ZQP 1ZQQ 1ZQS 1ZQT 1ZRC 1ZRD 1ZRE 1ZRF 1ZTG 1ZTT 1ZTW 1ZVV 1ZZI 1ZZJ 2A0I 2A3V 2A66 2AC0 2ACJ 2ADY 2AHI 2AJQ 2ATA 2AXY 2AYB 2AYG 2BDP 2BNZ 2BOP 2BPA 2BPF 2C5R 2C62 2C7O 2C7R 2C9L 2C9N 2CCZ 2CDM 2CGP 2D7D 2D7G 2D7H 2DGC 2DNJ 2DPD 2DPU 2DRP 2DWL 2DWM 2DWN 2E42 2E43 2EFW 2ES2 2ETW 2EUV 2EUW 2EUX 2EUZ 2EVF 2EVG 2EVH 2EVI 2EVJ 2EWJ 2EX5 2EZV 2FDC 2FIO 2FJV 2FJW 2FJX 2FKC 2FKH 2FL3 2FLC 2FLD 2FO1 2FVP 2FVR 2FVS 2G1P 2GE5 2GEQ 2GIE 2GIG 2GIH 2GII 2GIJ 2GLI 2GM4 2H1K 2H1O 2H27 2H7F 2H7G 2H7H 2H8C 2H8R 2HAP 2HAX 2HDD 2HHQ 2HHT 2HHU 2HHV 2HHX 2HMI 2HOS 2HOT 2HR1 2HT0 2HVR 2HVS 2HW3 2HZV 2I05 2I06 2I0Q 2I13 2I3P 2I3Q 2I5W 2I9K 2I9T 2IEF 2IIE 2IIF 2J6S 2J6T 2J6U 2KTQ 2KZM 2KZZ 2NL8 2NMV 2NRA 2NTC 2NTZ 2O49 2O4A 2O4I 2O61 2O6G 2O6M 2O93 2OA8 2OG0 2OR1 2OST 2OWO 2OXV 2OZB 2P0J 2P2R 2P5L 2P6R 2PI0 2PQU 2PRT 2PUB 2PUC 2PUD 2PUE 2PUF 2PUG 2PY5 2PYL 2PZS 2Q2K 2Q2T 2Q2U 2QBY 2QFJ 2QHB 2QL2 2QSG 2QSH 2R0Q 2R1J 2R2R 2R2S 2R2T 2R2U 2R5Y 2R5Z 2R9L 2RAM 2RBA 2RBF 2RGR 2RVE 2SSP 2UVR 2UVU 2UVV 2UVW 2UYC 2UYH 2UZ4 2V1U 2V2T 2V9W 2VA2 2VA3 2VHG 2VIC 2VIH 2VJU 2VJV 2VLA 2VOD 2VS8 2VW9 2VZ4 2W42 2W7N 2W7O 2W8K 2W8L 2W9B 2W9C 2WBS 2WBU 2WT7 2WTF 2WTY 2X6V 2XBM 2XE0 2XHB 2XM3 2XMA 2XO6 2XO7 2XQC 2XRO 2XRZ 2XSD 2YPA 2YPB 2YPF 2YVH 2Z3X 2Z6A 2Z6Q 2Z6U 2Z9O | 1679 |

| | | |
|---|---|---|
| | 2ZCJ 2ZKD 2ZKE 2ZKF 2ZO0 2ZO1 2ZO2 2ZZM 3A01 3A5T 3A5U 3AAF 3B39 3BAM 3BDP 3BI3 3BIE 3BJY 3BKZ 3BRD 3BRF 3BRG 3BS1 3BTX 3BTY 3BTZ 3BU0 3BUC 3C25 3C2I 3C2P 3C3L 3C46 3C58 3CBB 3CLC 3CLZ 3COQ 3CRO 3CVS 3CVT 3CW7 3CWA 3CWS 3CWT 3CWU 3D0A 3D0P 3D1N 3D4V 3D6Y 3D6Z 3D70 3D71 3DFV 3DFX 3DLB 3DLH 3DNV 3DO7 3DPG 3DSC 3DSD 3DVO 3DW9 3E3Y 3E40 3E41 3E42 3E43 3E44 3E45 3ECP 3EEO 3EH8 3EPH 3EPJ 3EPK 3ERE 3EXJ 3EXL 3EY1 3EYI 3EYZ 3F27 3F2B 3F2C 3F2D 3F73 3F8I 3F8J 3FD2 3FDE 3FDQ 3FHZ 3FMT 3FSI 3FTF 3G00 3G0Q 3G0R 3G2C 3G2D 3G38 3G3C 3G3Y 3G4T 3GA6 3GNA 3GNB 3GO8 3GP1 3GPP 3GPU 3GPX 3GPY 3GQ3 3GQ4 3GQ5 3GX4 3GXQ 3GYH 3H0D 3H15 3H25 3H8O 3H8R 3H8X 3HDD 3HJF 3HK2 3HM9 3HO1 3HOS 3HOT 3HPO 3HQF 3HTS 3HVR 3HXM 3HZI 3I2O 3I3M 3I49 3IAG 3IAY 3IGC 3IGK 3IGL 3IGM 3IKT 3IL2 3IV5 3JR9 3JRA 3JRB 3JRC 3JRD 3JRE 3JRF 3JRG 3JRH 3JRI 3JSO 3JSP 3JTG 3JXB 3JXC 3JXD 3K4X 3K57 3K58 3K59 3K9F 3KDE 3KET 3KJV 3KK1 3KK2 3KK3 3KMD 3KMP 3KO2 3KSA 3KSB 3KTQ 3KTU 3KXT 3KYL 3KZ8 3L2Q 3L2R 3L2U 3L2V 3L2W 3L4J 3L4K 3LDS 3LNQ 3LSR 3LTN 3LWH 3LWI 3M4A 3M9E 3M9M 3M9N 3M9O 3MAQ 3MDA 3MDC 3MDG 3MDI 3MFH 3MFI 3MFK 3MGH 3MGI 3MHT 3MIP 3MIS 3MKW 3MKY 3MKZ 3MVA 3MVB 3MX9 3MXB 3MXI 3MXM 3N1I 3N1J 3N1K 3N1L 3N4M 3N6S 3N7Q 3N97 3NGZ 3NH0 3NH1 3NH2 3NVI 3NVK 3O3I 3O6E 3O7V 3OA6 3ODA 3ODC 3ODE 3OGD 3OH6 3OH9 3OHA 3OHB 3OIN 3ON0 3OQM 3OQN 3OQO 3ORC 3OS0 3OS1 3OS2 3OSN 3OY9 3OYA 3OYB 3OYC 3OYD 3OYE 3OYF 3OYG 3OYH 3OYI 3OYJ 3OYK 3OYL 3OYM 3OYN 3P57 3P6Y 3PF4 3PKY 3PML 3PMN 3PNC 3POV 3PT6 3PVI 3PVP 3PVV 3Q05 3Q06 3Q0B 3Q0C 3Q0D 3Q0F 3Q1M 3Q2T 3Q2Y 3Q5F 3QFQ 3QLP 3QMB 3QMC 3QMD 3QMG 3QMH 3QMI 3QOQ 3QQY 3QRF 3QSV 3RZH 3RZJ 3RZK 3RZL 3RZM 3S3M 3S3N 3S3O 3S57 3S5A 3S6I 3S8Q 3SI8 3SJM 3SQI 3TAN 3TAP 3TAQ 3TAR 3TED 3TRZ 3TS0 3TS2 3TS8 3U2B 3U3Y 3U44 3U4Q 3U58 3U6C 3U6D 3U6E 3U6F 3U6L 3U6M 3U6O 3U6P 3U6Q 3U6S 3U6Y 3U7F 3U7G 3U7H 3UBT 3UBY 3UDG 3UFD 3UGO 3UGP 3UK3 3ULP 3UXW 3V1Z 3V20 3V21 3V6T 3V72 3V7J 3V7K 3VAG 3VAH 3VAI 3VAJ 3VAM 3VD0 3VD1 3VD6 3VDY 3VEK 3VKE 3VW3 3VW4 3VXV 3VXX 3VYB 3VYQ 3W3C 3WPC 3WPD 3WPE 3WPG 3WPH 3WPI 3WTS 3WTT 3WTU 3WTV 3WTW 3WTX 3WTY 3WU1 3ZKC 3ZLJ 3ZP5 3ZPL 3ZQL 3ZVK 4A04 4A75 4A76 4AA6 4AAB 4AAD 4AAE 4AAF 4AAG 4AIJ 4AIK 4AIL 4AQU 4AQX 4ATI 4ATK 4AUW 4AV1 4AWL 4B21 4B22 4B23 4B24 4BDP 4BDY 4BE1 4BNC 4CJA 4CRX 4CSA 4CYC 4D1Q 4D6N 4DAV 4DIH 4DII 4DK9 4DKJ 4DM0 4DQS 4DQY 4DWP 4E0D 4E0G 4E0J 4E0P 4E0Y 4E0Z 4E10 4E68 4E7H 4E7I 4E7J 4E7K 4E7L 4ENJ 4ENK 4ENM 4ENN 4ER8 4EUW 4F2J 4F6M 4F6N 4FGN 4FM9 4FXD 4G92 4H10 4HC7 4HC9 4HCA 4HJE 4IBU 4IBV 4IBW 4IUF 4IVZ 4IWR 4IX7 4IZZ 4J00 4J01 4J1J 4J2X 4JBK 4JBM 4JCX 4JCY 4KA4 4KFC 4KI2 4KNY 4KOE 4KPE 4KPF 4KTQ 4L0Y 4L0Z 4L18 4L5R 4L5S 4L8R 4LDX 4LG0 4LJR 4LNQ 4LOX 4LQ0 4LVI 4LVJ 4LVK 4LVL 4LVM 4LZ1 4LZ4 4LZG 4M04 4M0A 4M8O 4M94 4M95 4M9E 4MF2 4MGT 4MHT 4MKY 4MTD 4MTE 4MZR 4N41 4N47 4NDF 4NDG 4NDH 4NDI 4NHJ 4NM6 4NOE 4NW3 4OFA 4OFE 4OFH 4OI7 4OI8 4OMY 4ON0 4OO1 4OPJ 4OPK 4OQB 4OSH 4OSI 4OSJ 4OSK 4OSL 4OSM 4OSQ 4OSR 4OSS 4OST 4OSV 4OSW 4OSZ 4OT0 4OT3 4OTO 4OU6 4OU7 4OWW 4OWX 4P0P 4P0Q 4P9U 4PAR 4PBA 4PJO 4POG 4PSO 4PUQ 4PW5 4PW7 4PZI 4Q5V 4QCB 4QCL 4QEN 4QEO 4QEP 4QIL 4QPQ 4QQB 4QTJ 4QTK 4QTR 4R2A 4R2C 4R2D 4R2E 4R2P 4R2Q 4R2R 4R2S 4R55 4R56 4R79 4R8U 4RB1 4RB2 4RB3 4RBO 4RD5 4RDM 4RKG 4RKH 4RNM 4RNN 4RNO 4ROC 4ROD 4ROE 4RTK 4RTN 4RTQ 4RUA 4RUC 4RVE 4S0H 4S0N 4S2Q 4TNT 4TU8 4TU9 4TYN 4TZD 4UMK 4UN7 4UN8 4UN9 4UNA 4UNB 4UNC 4UNO 4UQM 4UT0 4UUS 4UUV 4UZB 4WK8 4Y60 4YEW 4YEX 4YEY 4YF0 4YFH 4YFT 4YGI 4Z3C 5BTF 5BTG 5BTI 5BTL 5BTN 5BYG 5CBX 5CBY 5CBZ 5CC0 5CC1 5CKY 5CL3 5CL4 5CL5 5CL6 5CL7 5CL8 5CL9 5CLA 5CLB 5CLC 5CLD 5CLE 5CO0 5CQQ 5CRJ 5CRK 5CRX 5CYS 5D23 5D2Q 5D2S 5D4R 5D4S 5D5U 5D5V 5D5W 5D5X 5D8F 5DLJ 5DNO 5DQT 5DQZ 5DS9 5DTD 5DTO 5DUI 5DWA 5DWB 5E17 5E18 5E24 5E3L 5E3M 5E3N 5E3O 5E69 5E6A 5E6B 5E6C 5E6D 5E8I 5ED4 5EEU 5EEV 5EEW 5EEX 5EEY 5EEZ 5EF0 5EF1 5EF2 5EF3 5EG6 5EIM 5EW1 5EYB 5F0Q 5F0S 5F55 5F56 5F8A 5FLV 5JUM 5T00 5T0U 5UND 5V3G 5VC8 5VC9 5W9Q 5W9S 5XFP 5XFQ 5XFR 5XS0 5YBB 5YBD 5YEF 5YEG 5YEH 5YEJ 5YEL 5YI3 5YIV 5YIW 5YJ3 5YUR 5YUS 5YUT 5YUU 5YUV 5YUW 5YUX 5YUY 5YUZ 5YV0 5YV1 5YV3 5YWS 5YWT 5YWU 5YWV 5YX2 5YYD 5YYE 5YZY 5YZZ 5Z00 5Z2T 5Z7I 5ZAD 5ZCW 5ZE0 5ZE1 5ZE2 5ZG9 5ZJQ 5ZJR 5ZJS 5ZJT 5ZK1 5ZKL 5ZKM 5ZKO 5ZLV 5ZMC 5ZMN 5ZMO 5ZQ0 5ZQ1 5ZQ8 5ZTH 5ZU1 5ZUO 5ZUP 5ZVA 5ZVB 5ZYU 5ZYV 6A2H 6A2I 6A57 6A8R 6AEG 6AKO 6AKP 6AMA 6AMK 6AS7 6B0O 6B0P 6B0Q 6B0R 6B1Q 6B1R 6B1S 6BEK 6BHX 6DKS 6KBZ 6KC7 6KC8 6KCQ 6KCS 6KDV 6KI3 6KIJ 6KKS 6KVO 6L6S 6L6Y 6L84 6LAE 6LAU 6LBI 6LC1 6LEW 6LRD 6LTY 6LWA 6LWB 6LWC 6LWD 6LWF 6LWG 6LWH 6LWI 6LWJ 6LWK 6LWL 6LWM 6LWN 6LWO 6LWP 6LWR 6LXN 6M75 6MIG 6MIH 6MIK 6ML2 6ML3 6ML4 6ML5 6ML6 6ML7 6MP3 6MQ8 6MR7 6MR8 6MRJ 6MXO 6NCE 6NCM 6NJQ 6NSN 6NSR 6NUA 6NUH 6O19 6O8F 6O8H 6R2V 6SKO 6T78 6U8P 6U8V 6U8W 6U8X 6U90 6U91 7C4Q 7C4R 7C98 7C99 7C9A 7C9C 7CC9 7CCD 7CCJ 7CSZ 7CUH 7D3T 7DCI 7DCJ 7DCS 7DCT 7DCU 7DW5 7EF8 7EF9 7ET4 7ET6 7EYI 7ICE 7ICF 7ICG 7ICH 7ICI 7ICJ 7ICK 7ICL 7ICM 7ICN 7ICO 7ICP 7ICQ 7ICR 7ICS 7ICT 7ICU 7ICV 7JSA 7JZ0 | |
| Total | | 1837 |

| **Table S2.** List of PDB IDs used for benchmarking the optimal cut-off distance between Cα — O5′ |
|---|
| 2FJ7, 7KTQ, 4QLC, 7ON1, 4X23, 7PII, 6MUP, 7VDT, 7KBD, 7XD1 |

**Table S3.** The computational time required to complete the task for 10 representative PDB's

| **Task** | **Time (MM:SS)** | **Total time (MM:SS)** |
|---|---|---|
| **Residue pair distance (CA-O5')** | **73:18** | |
| **Heavy atom-to-atom contacts for qualifier residue pairs at distinct cut-off (nm)** | | |
| 0.9 | 73:18 + 09:11 | 82:29 |
| 1.0 | 73:18 + 13:26 | 86:44 |
| 1.1 | 73:18 + 28:25 | 101:43 |
| 1.2 | 73:18 + 45:26 | 118:44 |
| 1.3 | 73:18 + 71:40 | 144:58 |
| **1.2 (for R and K) and 1.0 for others** | **73:18 + 24:01** | **97:19** |
| **Common Method (All heavy atom-to-atom method)** | | **37080:01** |

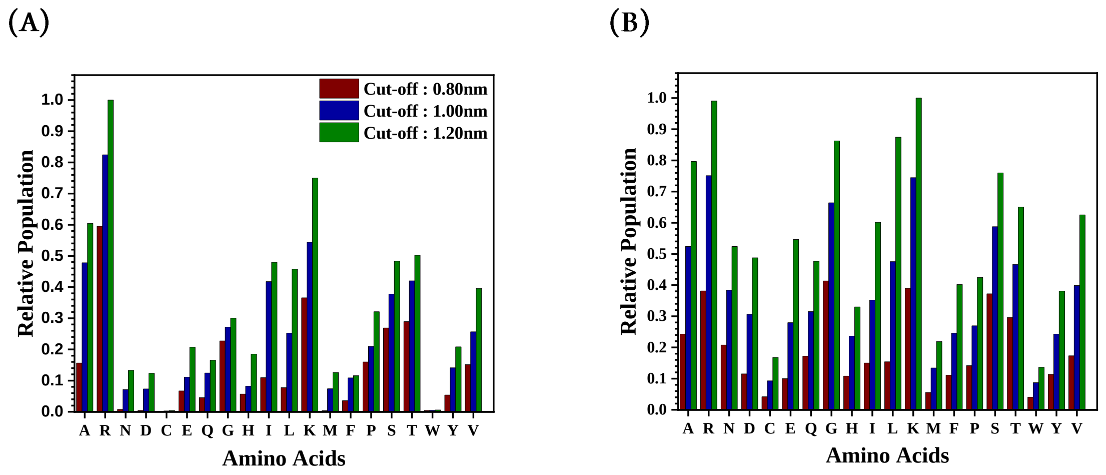


**Figure S1.** The relative population of all the 20 standard amino acids involved at the interface with 0.8 to 1.2nm cut-off distances between Cα and O5′ for (A) nucleosomes and (B) non-nucleosomes. The number PDB structures considered, with better than 3.5 Å resolution, for nucleosomes and non-nucleosomes were 258 and 1679, respectively. The legends same for both graphs.
